# Enhanced polar auxin transport in tomato polycotyledon mutant seems to be related to glutathione levels

**DOI:** 10.1101/656249

**Authors:** Sapana Nongmaithem, Rachana Ponukumatla, Yellamaraju Sreelakshmi, Pierre Frasse, Mondher Bouzayen, Rameshwar Sharma

## Abstract

Glutathione-dependent root growth in Arabidopsis is linked to polar auxin transport (PAT). Arabidopsis mutants with reduced glutathione (GSH) levels also show reduced PAT. To gain an insight into the relationship between PAT and GSH level, we analysed tomato polycotyledon mutant, *pct1-2*, which has enhanced PAT. Microarray analysis of gene expression in *pct1-2* mutant revealed underexpression of several genes related to glutamate and glutathione metabolism. In consonance with microarray analysis, enzymatic as well as *in-vivo* assay revealed higher glutathione level in the early phase of *pct1-2* seedling growth than WT. The inhibition of auxin transport by 2,3,5-triiodobenzoic acid (TIBA) reduced both GSH level and PIN1 expression in *pct1-2* root tips. The reduction of *in vivo* GSH accumulation in *pct1-2* root tips by buthionine sulfoximine (BSO) stimulated elongation of the short root of *pct1-2* mutant akin to TIBA. The rescue of short root phenotype of *pct1-2* mutant was restricted to TIBA and BSO. The other auxin transport inhibitors 1-*N*-naphthylphthalamic acid (NPA), 2-[4-(diethylamino)-2-hydroxybenzoyl] benzoic acid (BUM), 3-chloro-4-hydroxyphenylacetic acid (CHPAA), brefeldin and gravacin inhibited root elongation in both WT and *pct1-2* mutant. Our results indicate a relationship between PAT and GSH level in tomato akin to Arabidopsis. Our work also highlights that TIBA rescues short root phenotype of the *pct1-2* mutant by acting on a PAT component distinct from the site of action of other PAT inhibitors.

## Introduction

In nature, plant growth involves continuous adaptation to its environment by integration and coordination of signaling pathways within a tissue or different organs. Above integration is achieved by the transfer of information between different organs using plant hormones, small metabolites, and peptides. Among the plant hormones, auxin [Indole acetic acid (IAA)] plays a major role enabling short-term responses to environment such geotropism and phototropism, of long-term adaptation to environment, such as onset of dormancy or determination of plant architecture (Zažímalová et al. 2014; Vanneste and Friml 2009; Davies 2004)). Besides being a master regulator of a plethora of plant developmental responses, auxin also plays an important role in modulating plant growth responses to different abiotic or biotic stresses (Kazan and Manners 2009; Cheyong et al. 2002). A major interaction of auxin with environmental signal involves the modulation of redox signaling that may involve cellular homeostasis of reactive oxygen species (ROS) and induction of enzymes leading to ROS detoxification (Iglesias et al. 2010; Tognetti et al. 2012).

The directional mobility of auxin within the plants is key to the regulation of several developmental responses. The directional movement of auxin is carried out by plasma membrane-localized transporters, which facilitate auxin influx and efflux from neighboring cells (Chen and Baluška 2013). The auxin transporters comprise of diverse groups consisting of AUX/LAX proteins carrying out auxin influx (Marchant et al. 1999), auxin efflux by members of PIN-FORMED (PIN) family (Petrásek and Friml 2009), and the P-glycoprotein/multidrug resistance B family of ATP-binding cassette transporter B-type (ABCB) proteins (Wu et al. 2007). The role of these auxin transporters in plant development is largely inferred from analysis of mutations. The mutations in AUX/LAX protein lead to impaired gravitropic responses ((Marchant et al. 1999), in PIN1 protein causes naked inflorescence in Arabidopsis (Okada et al. 1991), and alleles of PGP proteins alters the plant architecture in maize leading to dwarf plants (Multani et al. 2003). ROS production under stress also influences auxin transport by affecting the activity of auxin transporters. The environmental stresses and biotic stresses can alter the cellular auxin levels by influencing the polar localization of PIN proteins (Grunewald and Friml 2010).

The interaction between auxin and ROS is best exemplified in studies on root growth and elongation. The cysteine-containing tripeptide glutathione (GSH) and ascorbate (ASC) seems to play an important role in the regulation of root apical meristem (De Tullio et al. 2010). In maize root tips, auxin maxima in quiescent center cells are associated with lower levels of ASC and GSH (Jiang et al. 2003). In Arabidopsis, ASC and GSH affect root length and mitotic activity (Sanchez-Fernandez et al. 1997). The mutations in the ROOT MERISTEMLESS1 (*RML1*) gene, which encodes γ-glutamylcysteine synthetase (γ-ECS) gene, regulating the first step of GSH biosynthesis, causes defects in root apical meristem and reduced the formation of post-embryonic roots (Vernoux et al. 2000). A relationship between GSH level and auxin transport was shown by depleting endogenous GSH level by buthionine sulphoximine (BSO), an inhibitor of γ-ECS (Koprivova et al. 2010). The BSO-mediated reduction of Arabidopsis root growth was also associated with the reduced expression of PIN proteins. In *cad2* mutant, that is allelic to *rml1* mutant (Vernoux et al. 2000), polar auxin transport (PAT) was reduced to 10% of wild type level, indicating a relationship between glutathione level and PAT (Bashandy et al. 2010). Likewise, *ntra ntrb cad2* triple mutant of Arabidopsis (Bashandy et al. 2010), bearing mutations in NADPH-dependent thioredoxin reductases and γ-glutamylcysteine synthetase shows naked inflorescence like *pin1* (Gälweiler et al. 1998) and *pinoid* (*pid*) (Bennett et al. 1995) mutants. The above mutant also shows reduced auxin transport, and its phenotype of reduced flowers on inflorescence can be reversed by exogenous GSH application.

These mutant analyses indicated that GSH plays an important role in regulating root growth and development and also interacts with PAT. In contrast to Arabidopsis *ntra ntrb cad2* mutant, tomato *pct1-2* mutant shows enhanced PAT and increased number of flowers in the inflorescence (Al-Hammadi et al. 2003). The diametrically opposing phenotypes of *ntra ntrb cad2* and *pct1-2* mutants prompted us to examine whether there is a relationship between GSH level and auxin transport in the *pct1-2* mutant. Microarray analysis was used to ascertain, whether the *pct1-2* mutation affects the expression of glutathione-related genes. We show that *pct*1-2 mutant exhibits enhanced GSH levels during early seedling development. The similarity between BSO- and TIBA-mediated rescue of short root phenotype of *pct*1-2 mutant indicates a linkage between PAT and GSH levels in tomato seedlings.

## Results

The striking enhancement in the polar auxin transport (PAT) in tomato *pct1-2* mutant is associated with several developmental abnormalities throughout the life cycle. Light-grown *pct1-2* mutant seedlings display shorter root and smaller epidermal cells in hypocotyls and cotyledon than wild type. The *pct1-2* hypocotyls also show alteration in anatomy with the abnormal placement of vascular bundles and reduction of the central pith (Al-Hammadi et al. 2003). Considering these development abnormalities, the gene expression profiles of the *pct1-2* mutant was compared with wild type in roots, hypocotyls, and cotyledons.

### Genes differentially expressed in *pct*1-2 seedlings

In conformity with diverse development milieu of organs, the differentially expressed genes belonging to different functional categories revealed organ-specific patterns. All functional categories combined, the differential expression of the genes was more skewed towards down-regulation in *pct1-2* mutant rather than upregulation (Cotyledon 92↓, 55↑; hypocotyl 81↓, 31↑; root 118↓, 55↑) (Figure 1, **Supplementary Table 1**). Excluding photosynthesis genes in roots, in all three organs, the genes belonging to the stress category displayed the highest downregulation as well as upregulation. An examination of differentially underexpressed genes in *pct1-2* mutant revealed that several of these regulated the GSH metabolism (**Supplementary Table 2**). These belonged to three categories viz. lactoylGSH lyase family (↓ root, hypocotyls, and cotyledons), glutatredoxin genes (↓ hypocotyls), and glutathione-S-transferases (↓ hypocotyls and roots, ↑ cotyledons). Each of these groups plays unique roles in the plant metabolism and yet strongly associate with glutathione, albeit in a different way. Assuming that the differential expression of these genes reflects the alteration in endogenous GSH level, we compared the GSH levels in *pct1-*2 with wild type seedlings.

**Figure 1:**
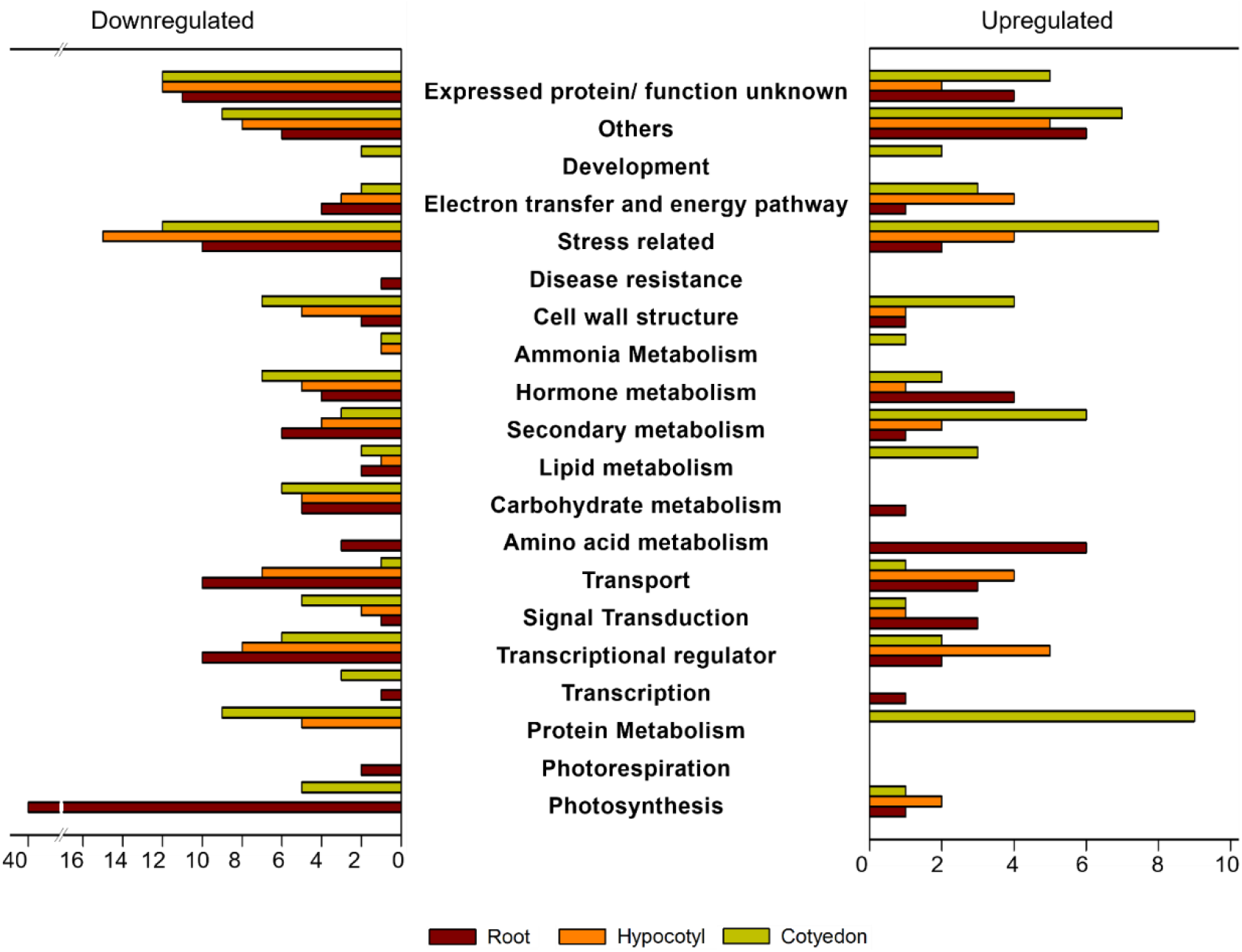
Functional grouping of differentially expressed genes in cotyledons, roots and hypocotyls of 5-day old *pct1-2* mutant seedlings. For differential expression, only genes with ≥1.4 fold changes were considered.

### *pct1-2* seedlings have more GSH in early stages of development

The analysis of GSH levels in the seeds and developing light-grown seedlings revealed that *pct*1-2 mutant had a higher level of GSH in dry seeds than WT. Post-germination at 2 days, GSH level increased by 3 fold in *pct*1-2 and 2.75 fold in WT seedling, followed by a decline in GSH level in both pct1-2 mutant and WT seedlings with nearly similar levels in 7-day old seedlings (Figure 2A). The estimation of GSH levels in individual organs of the 5-day old seedlings (at what stage, light dark, age) revealed that while the GSH levels were similar in cotyledons and hypocotyls, it was considerably higher in roots of *pct*1-2 mutant (Figure 2B). Considering that the roots of the *pct*1-2 mutant are shorter than the wild type, the higher GSH level in roots may have some association with the above phenotype. The staining root tips with monochlorobimane for *in vivo* GSH level showed more uniform stain in *pct1-2* mutant root tips compared to WT where it was localised in epidermal layers (Figure 3A). The treatment of auxin transport inhibitor TIBA that stimulate the root elongation in the *pct1-2* mutant (Al-Hammadi et al. 2003) shifted monochlorobimane staining pattern in *pct1-2* root tips akin to wild type control. Immunochemically localization of PIN1 protein in root tips revealed that TIBA also reduced the PIN1 localization in both WT and *pct1-2* root (Figure 3B).

**Figure 2:**
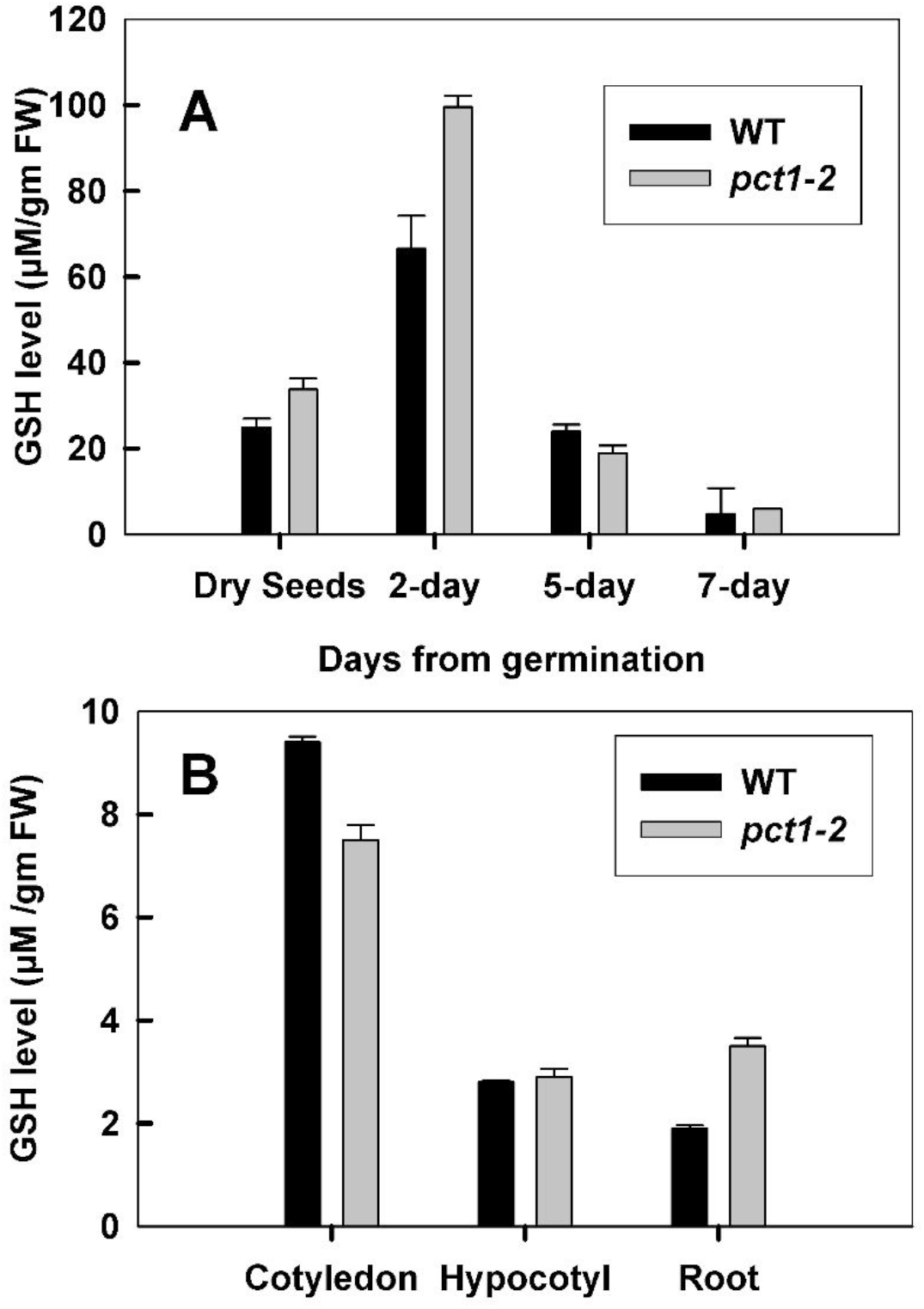
The levels of GSH in the tomato WT and *pct1-2* mutant seedlings. **A.** GSH level at different stages of seedling development. **B.** GSH level in different organs of 5-day old seedling.

**Figure 3:**
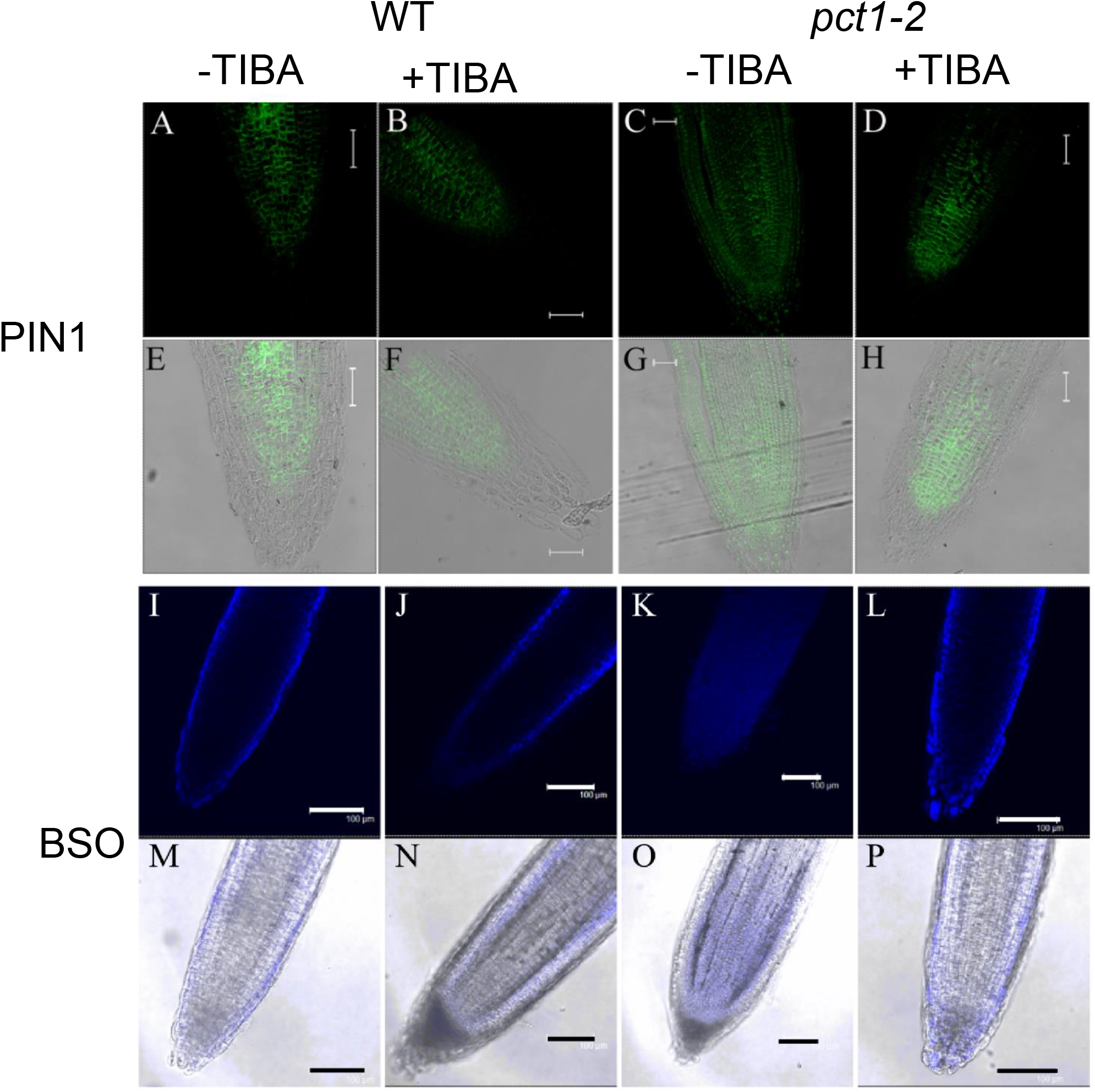
*In situ* imaging of PIN1 (A-H) and GSH (I-P) localization in root tips excised from 5-day old light-grown WT and *pct1-2* seedlings. The seedlings were grown in the presence or absence of 0.5 µM TIBA. The data represents the average of at least three independent biological replicates. Scale - 100 µM.

### Light-grown WT seedlings were more sensitive to GSH than *pct*1-2

Considering that WT and *pct1-2* seedlings had different GSH level, we next examined whether exogenous GSH influences tomato seedling growth. The responsiveness of light-grown *pct1-2* seedlings to GSH considerably differed from WT (Figure 4A,B). Light-grown *pct1-2* seedlings responded in biphasic fashion, wherein in the first phase the lower GSH concentration inhibited *pct1-2* hypocotyl and root growth, similar to WT seedlings. However, at higher GSH concentrations (> 1.5 mM) *pct1-2* hypocotyls did not show any further decrease in length and were taller than WT. Likewise, *pct1-2* roots were resistant to higher GSH concentration (>1 mM), and at 2.5 mM concentration, the *pct1-2* roots were distinctly longer than WT. The *pct1-2* resistance to GSH was restricted to light-grown seedlings; the dark-grown *pct1-2* seedlings showed growth inhibition similar to WT (Figure 4C,D). However, the responsiveness to growth inhibition in dark-grown seedlings was observed only at GSH concentration higher than 0.5 mM and 1.5 mM for hypocotyl and roots, respectively.

**Figure 4:**
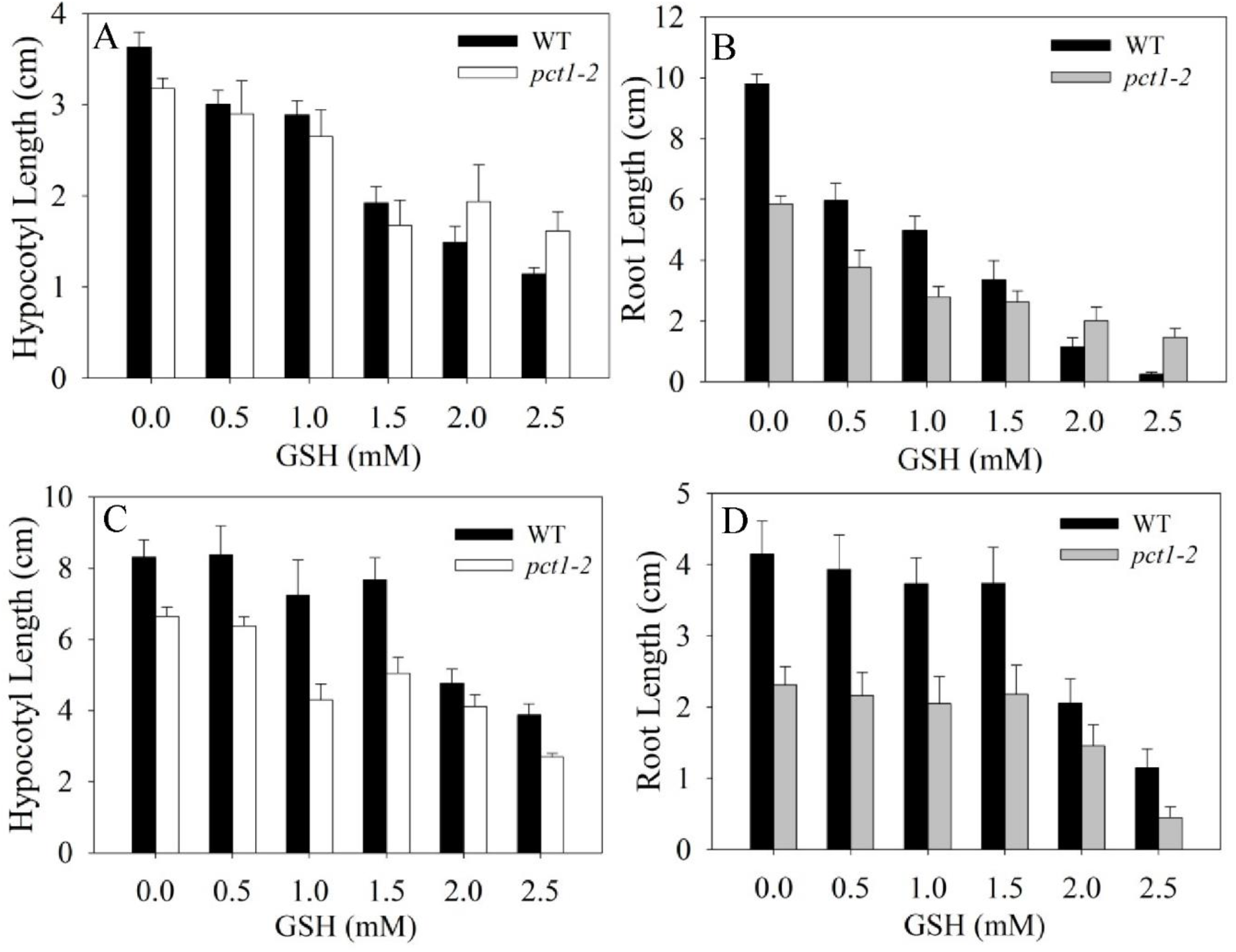
Effect of GSH on the hypocotyl (A,C) and root (B,D) growth of light-(A,B) and dark-grown (C,D) 5-old tomato WT and *pct1-2* mutant seedlings. Values are the means ± SE from five independent replicates.

### BSO differentially affects root length in WT and *pct*1-2 seedlings

To decipher the relationship between GSH and root/hypocotyl growth, we examined whether the depletion of GSH differentially affects above growth responses. For this, we used buthionine sulfoximine (BSO) which specifically inhibits γ-glutamylcysteine synthetase involved in the first step of GSH synthesis. The increasing concentrations of BSO mildly inhibited elongation of pct1*-2* hypocotyls and did not affect WT. In contrast, *pct*1-2 showed stimulation of root elongation at 0.2 mM BSO, followed by reduced stimulation up to 0.6 mM BSO (Figure 5A,B). Contrastingly, BSO inhibited WT root elongation at higher than 0.2 mM concentration. The observed elongation of the *pct1-2* root at a lower dosage of BSO was contrary to the report that BSO inhibits root elongation in WT and auxin-resistant mutants of Arabidopsis (Koprivova et al. 2010).

**Figure 5:**
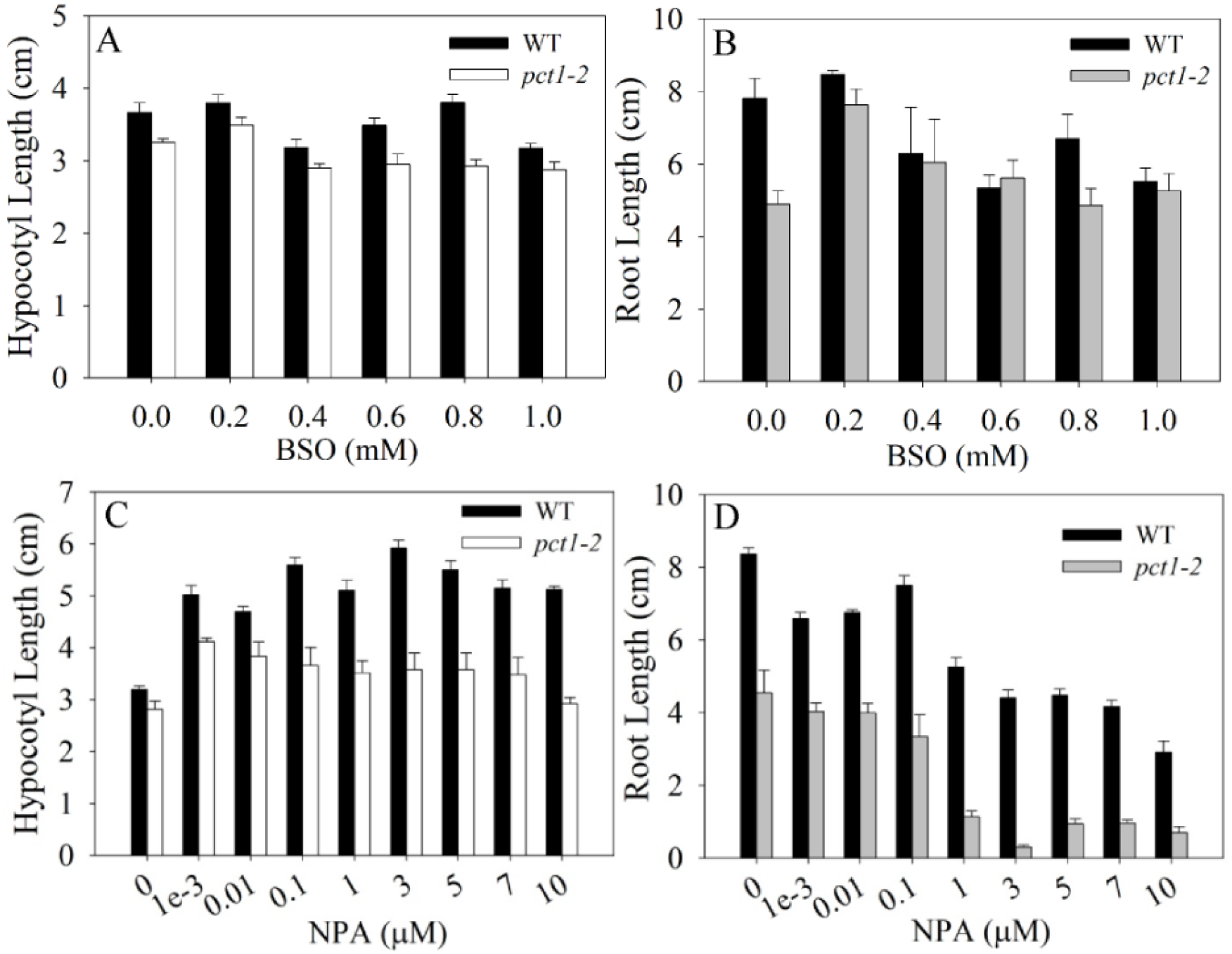
Effect of BSO (A,B) and NPA (C,D) on the hypocotyl (A,C) and root (B,D) growth of light-grown 5-old tomato WT and *pct1-2* mutant seedlings. Values are the means ± SE from five independent replicates.

### Is enhanced PAT in *pct*1-2 due to the increased shuttling of PIN proteins?

The enhanced auxin transport in tomato *pct1-2* mutant is associated with enhanced PIN1 protein localization in root tips (Kharshiing et al. 2010). Considering that BSO triggers loss of PIN1 proteins in primary roots of Arabidopsis (Koprivova et al. 2010), we examined whether the observed root elongation in the *pct*1-2 mutant due to influences of BSO on intracellular shuttling of PIN proteins. To ascertain this, we used brefeldin A (BFA), a fungal toxin, which inhibits vesicle transport resulting in intracellular accumulation of PIN1 protein. Both WT and *pct1-2* mutant seedlings grown in the presence of BFA, showed biphasic growth inhibition of root elongation (Figure 5C,D). The BFA mildly reduced the root elongation up to 0.25 µM, followed by sharp inhibition of root elongation at higher concentration. Considering both WT and *pct*1-2 responded similarly to BFA, it appears that BSO-mediated root elongation in the *pct1-2* mutant was not related to PIN1 trafficking.

### Do other auxin transporters play a role in enhanced PAT in *pct*1-2?

We examined whether other inhibitors of auxin transport phenocopy the responses elicited by glutathione and BSO on the *pct1-2* mutant. Since *pct1*-2 display the enhanced PAT, and reduction of PAT increases the root elongation, the reduction in auxin influx mediated by AUX1 may have a similar effect. The auxin influx mediated by the AUX1 protein is considerably inhibited by **CHPAA** (Parry et al. 2001). CHPAA differentially inhibited hypocotyl elongation wherein *pct1-2* mutant was more sensitive to growth inhibition than WT (Figure 6A,B). Contrarily CHPAA stimulated elongation of WT roots at lower concentration (<15 µM), while *pct1-2* mutant root showed growth inhibition at 5 µM followed by a further reduction in growth at a higher dosage. Interestingly N-1-naphthylphthalamic acid **(NPA)** that targets ABCB transporters induced hypocotyl elongation in both WT and *pct*1-2 seedlings (Teale and Palme 2017) (Figure 6C,D). Oppositely, NPA inhibited root elongation in both WT and *pct1*-2, in biphasic fashion with higher inhibition elicited at concentration > 1 µM. The inhibition of ABCB1, an auxin transporter, with **BUM** (2-(4-Diethylamino-2-hydroxybenzoyl) benzoic acid) (Kim et al. 2010) elicited mild enhancement in hypocotyl and root elongation in *pct1-2* seedlings followed by growth inhibition at higher concentration (> 10 µM) (Figure 7A,B). While BUM did not inhibit WT hypocotyl elongation, it inhibited root elongation Gravacin, that specifically inhibits PGP19 auxin transporter (Rojas-Pierce et al. 2007) did not inhibit hypocotyl elongation but inhibited root elongation both in WT and *pct*1-2 (Figure 7C,D).

**Figure 6:**
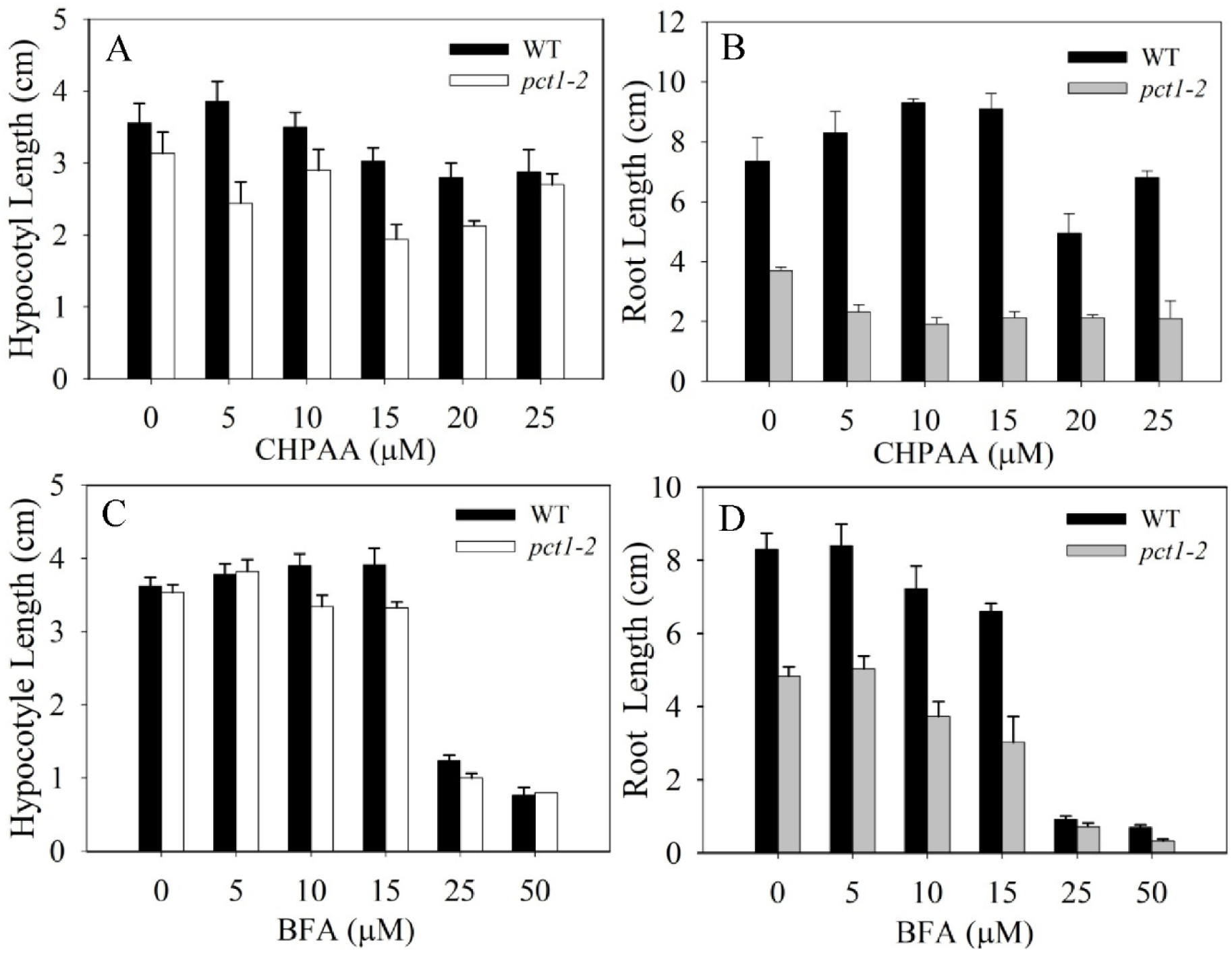
Effect of CHPAA (A,B) and BFA (C,D) on the hypocotyl (A,C) and root (B,D) growth of f light-grown 5-old tomato WT and *pct1-2* mutant seedlings. Values are the means ± SE from five independent replicates.

**Figure 7:**
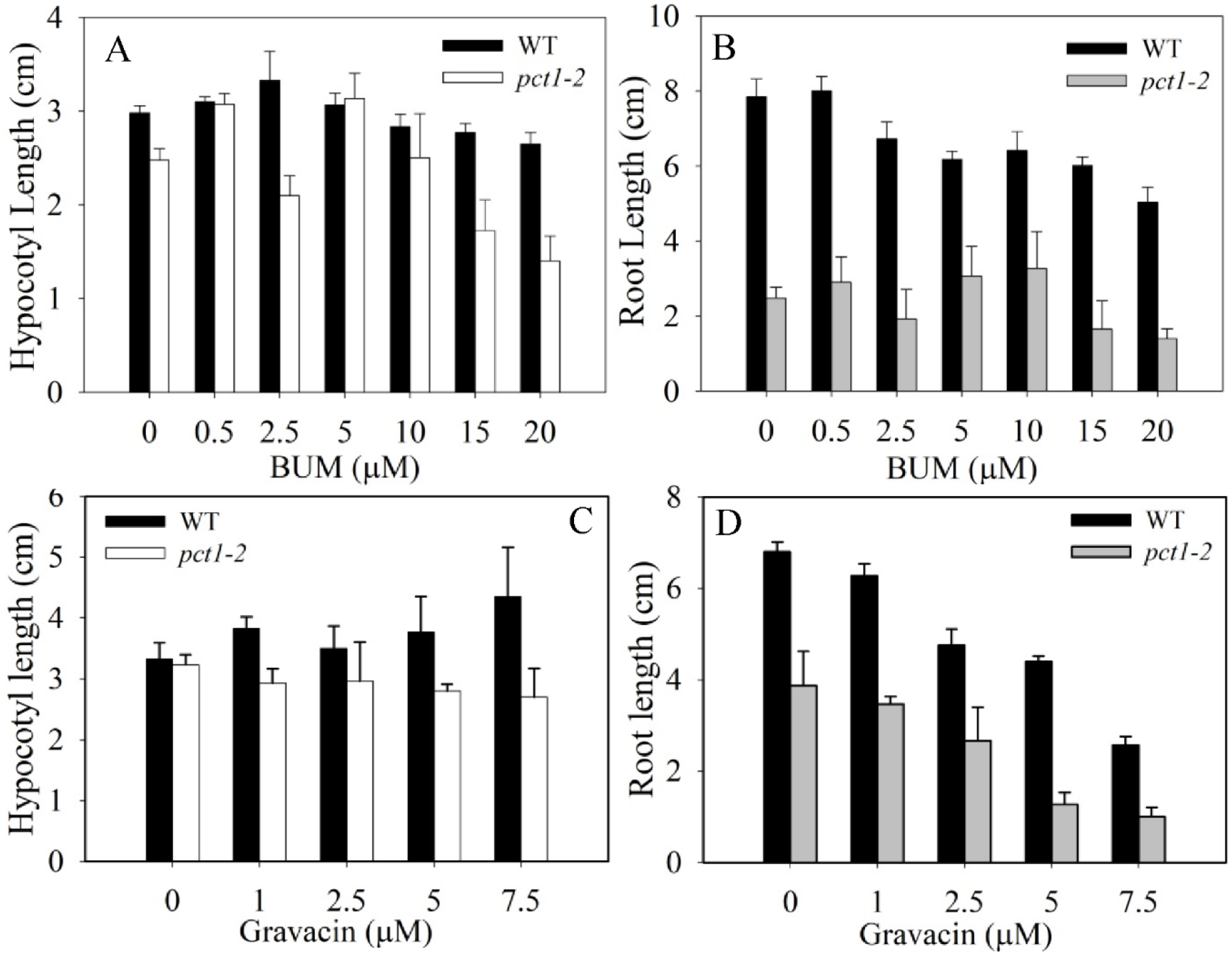
Effect of BUM (A,B) and gravacin (C,D) on the hypocotyl (A,C) and root (B,D) growth of light-grown 5-day old tomato WT and *pct1-2* mutant seedlings. Values are the means ± SE from five independent replicates

## Discussion

Our study examined two aspects; first, whether there is a potential link between glutathione and enhanced auxin transport in the *pct1*-2 mutant. Second, can the shorter root phenotype of *pct1*-2 mutant be rescued by other PAT inhibitors, as was observed for TIBA (Al-Hammadi et al. 2003). In Arabidopsis, *ntra ntrb cad2* mutant produces naked inflorescence stem, reminiscent of *PIN1* mutant that has reduced auxin transport (Bashandy et al. 2010). Consistent with naked inflorescence stem, above mutant also show highly reduced PAT and reduced abundance of PIN1 protein. The GSH level in *ntra ntrb cad2* mutant is 32% of WT, and application of GSH partially rescues the mutant phenotype. Collectively, the above data indicate a relationship between endogenous GSH levels and auxin transport in Arabidopsis.

Contrasting to Arabidopsis *ntra ntrb cad2* mutant, tomato *pct1-2* mutant has enhanced PAT (Al-Hammadi et al. 2003); shows overexpression of PIN1 protein (Kharshiing et al. 2010). Therefore, it can be assumed that the enhanced PAT in the *pct1-2* mutant may have a relationship with endogenous GSH levels. This assumption is reinforced by the observation that the majority of genes involved in GSH metabolism are underexpressed in all three organs of *pct1-2* seedlings relative to WT. The *pct1-2* seedlings show elevated GSH level in dry seeds and during early seedling development. The *pct1-2* seedlings have shorter roots and show higher GSH level in roots. In particular, *pct1-2* root tip has higher *in situ* GSH localization than WT. Notably, the application of TIBA that stimulates *pct1*-2 root elongation shifts GSH staining pattern akin to WT. Broadly, these results point to a relationship between GSH and PAT in tomato too.

The application of increasing dosage of GSH revealed the differential influence of GSH on light-grown seedling growth of WT and *pct1-2.* In tomato, the elongation of adventitious roots is dependent on optimal GSH level in the root tip, and exogenous auxin possibly inhibits root growth by the accumulation of supra-optimal GSH levels (Tyburski and Tretyn 2010). Consistent with this, at a lower dosage, GSH inhibited root/hypocotyl growth of WT and *pct1-2* seedlings. However, at a higher dosage of GSH, light-grown *pct1-2* seedlings showed resistance to the GSH-induced growth inhibition of root and hypocotyls. Earlier studies have shown that exogenous GSH rescues the phenotypes of Arabidopsis mutants (Li et al. 2006, Bashandy et al. 2010). Expectedly, exogenous GSH exacerbated *pct1*-2 shorter root phenotype. However, at higher GSH dosage resistance of *pct1*-2 may due to either reduced uptake of GSH or attainment of peak inhibition level elicited by GSH. Since dark-grown *pct1*-2 and WT seedling show nearly similar GSH-induced growth inhibition, it appears that resistance to a higher dosage of GSH was a feature specific to light-grown *pct1*-2 seedlings.

In Arabidopsis, application of buthionine sulfoximine (BSO) which specifically inhibits γ-glutamylcysteine synthetase, an enzyme involved in GSH biosynthesis, leads to a reduction in the root growth due to GSH depletion (Koprivova et al. 2010). The BSO-triggered GSH depletion is also associated with loss of PIN1 expression, which likely leads to reduced PAT (Koprivova et al. 2010; Bashandy et al. 2010). Considering that *in vivo* reduction of GSH reduces PAT, it can be assumed that the application of BSO should rescue short root phenotype of *pct1*-2, akin to TIBA (Al-Hammadi et al. 2003). Consistent with this assumption, 0.2 mM BSO stimulated *pct1*-2 root elongation by 47% akin to TIBA, whereas it had only a mild effect on WT roots. The inefficacy of higher than 0.2 mM BSO dosage on *pct1*-2 root elongation may be due to interference with the role of GSH by BSO in other cellular responses. Taken together with high GSH level in the *pct1-2* root, differential expression of GSH-related genes, growth inhibition by GSH and BSO-mediated *pct1*-2 root rescue seem to indicate that *pct1*-2 mediated alteration in cellular homeostasis of GSH level may have a relationship at least with short root phenotype of the mutant. In *ntra ntrb cad2* mutant application of GSH induces flower formation on the naked inflorescence. In contrast, *pct1*-2 mutant produces a very high number of flowers in the inflorescence (Al-Hammadi et al. 2003). It is likely that enhanced flower phenotype of *pct1*-2 may be related to GSH level. Nonetheless, the above observation indicated a putative linkage between endogenous GSH level and enhanced PAT in *pct1*-2.

Even though our study, as well as earlier studies (Bashandy et al. 2010; Koprivova et al. 2010), indicates a relationship between endogenous GSH level and auxin transport, the linkage between these two processes is indirect and may involve other cellular intermediates. Similar to GSH deficient mutant, flavonoids deficient Arabidopsis mutants show altered expression and localization PIN proteins (Peer et al. 2004). The flavonoid deficient Medicago plants also show enhanced auxin transport (Wasson et al. 2006). However, the *pct1*-2 mutant plants have similar flavonoids levels at least in hypocotyls to WT, thus discounting the possibility of the role of flavonoids in enhanced PAT (Al-Hammadi et al. 2003).

The similarity between BSO-mediated and TIBA-mediated rescue of *pct1*-2 root elongation entails that both may affect a similar subset of cellular responses. The TIBA application to plants also induces glutathione S-transferases (GSTs) (Flury et al. 1995, 1996). The increased level of GSTs can reduce intracellular GSH concentration by its conjugation to endogenous electrophilic compounds (Chronopoulou et al. 2017). The microarray studies indicated that GST transcripts are underexpressed in the roots and hypocotyl of *pct1-2* mutant. The reduced GST expression may have contributed to increased GSH levels in *pct1-2* roots. TIBA is also known to react non-enzymatically at least *in vitro* with sulfhydryl compounds such as cysteine, GSH, and coenzyme A (Leopold and Price 1957). Such an interaction between GSH and TIBA *in vivo* has the possibility of reducing intracellular concentration of GSH. The *in situ* staining of GSH levels in WT and *pct1-2* root supports the possibility of such an interaction. Compared to WT, *pct1-2* root shows a higher level of GSH as visualized by MCB staining. Importantly, the incubation with 0.5 µM TIBA, which stimulation elongation of *pct1-2* roots (Al-Hammadi et al. 2003) also reduces the GSH staining in mutant root tips. Similarly, TIBA also reduces the PIN1 protein level in WT and *pct1-2* root tips. These results indicate a two-prong action of TIBA in tomato root tips involving in the parallel reduction of GSH levels and PIN1 level in root tips. Similar to TIBA, BSO reduces PIN1 localization (Koprivova et al. 2010; Bashandy et al. 2010) as well as GSH levels (Zhu et al. 2016) in roots. Taken together, the observed reduction in GSH level in *pct1-2* roots may result from either TIBA-mediated induction of GSTs or direct interaction of TIBA with endogenous GSH, or a combination of both.

The similarity between TIBA and BSO action, at least in the reduction of PIN1 localization and GSH levels, indicates that though their primary action may be different, they may be converging at a similar subset of responses regulating auxin transport. This convergence seems at the GSH level, as both reduce GSH level in root tips. While the reduction in GSH level in *ntra ntrb cad2* mutant is associated with reduced PAT, a higher rate of PAT in the *pct1-2* mutant is associated with higher GSH levels. These evidences suggest that higher level of GSH in the *pct1-2* root has a linkage with PAT. Taking cognizance that BSO and TIBA likely rescue short root phenotype by affecting the PAT, we examined whether other reported auxin transport inhibitors can rescue the short root phenotype of *pct1-2* mutant.

Among the PAT inhibitors, 1-N-naphthylphthalamic acid (NPA) is the most widely used to auxin-related developmental response such as phyllotaxy (Reinhardt et al. 2000) and embryo development (Schiavone and Cooke 1987). The NPA and TIBA seem to act on different subsets of the auxin transport system. The *in vitro* studies indicated that TIBA acts directly on some component of PAT, as it reversibly competes with auxins, whereas NPA inhibited PAT by binding to a site different from auxin binding site (Thomson et al. 1973). Unlike NPA, radiolabelled TIBA also shows the basipetal polar transport similar to auxin in corn coleoptiles (Thomson et al. 1973). The action of NPA on PAT is attributed to its interaction with several cellular components, such as twisted-dwarf-1 (TWD1), ABCB1, ABCB19 (Noh et al. 2001), plasma membrane-associated aminopeptidases APM1 and APP1 (Murphy et al. 2002). It also alters the intracellular cycling of PIN proteins between endosomal vesicles and the plasma membrane (Geisler et al. 2005).

In contrast to TIBA which rescued *pct1-2* root elongation, both WT and *pct1-2* mutant seedling grown on varying NPA concentration display reduction in root elongation. Surprisingly, NPA stimulated hypocotyl length elongation in both WT and *pct*1-2, with the effect being milder in *pct1-2*. The NPA-mediated hypocotyl elongation in tomato was also observed by Kraepiel et al. (2001). The opposite effect of NPA and TIBA on *pct1-2* mutant root elongation indicates that these two inhibitor influences a different subset of PAT. Moreover, only the PAT subset affected by TIBA and BSO is able to stimulate the root elongation in the *pct1-2* mutant.

The dichotomy between NPA and TIBA action indicated that TIBA rescued *pct1-2* root elongation by acting on a unique subset of PAT that is also influenced by GSH. In such a scenario, other known auxin transport inhibitors should also be ineffective in rescuing *pct1-2* root elongation. Treatment of Arabidopsis roots with brefeldin (BFA) prevents intracellular cycling of PIN1, leading to accumulation of PIN1 in the endodermal compartment (Geldner et al. 2001). BFA inhibited both root and shoot elongation of WT and *pct1-2* mutant. Interestingly, auxin influx inhibitor, CHPAA (Parry et al. 2001) stimulated elongation of WT roots at a lower dosage. However, CHPPA like BFA inhibited WT and *pct1-2* mutant hypocotyl elongation. Likewise, BUM (Kim et al. 2010), an ABCB inhibitor, did not elicit any significant rescue of the *pct1-2* mutant root, rather inhibited it at a higher dosage. Similarly, gravacin (Rojas-Pierce et al. 2010), a PGP19 auxin transporter inhibitor also did not elicit any rescue of the *pct1-2* root, rather inhibited both hypocotyl and root elongation. Taken together, these results reinforce the view that the TIBA and BSO affect a different subset of auxin transport affected by the *pct1-2* mutation. This subset involves a PAT component, which is modulated only by TIBA and BSO. The other known auxin transport inhibitors do not interact with this component at least in *pct1-2* mutant seedlings.

It remains to be determined how *pct1-2* mutation influences two diverse response such as overexpression of PIN1 protein (Kharshiing et al. 2010) and increase in GSH level. In case higher GSH level is the primary action of *pct1-2* mutation, how higher GSH level leads to overexpression of PIN1 protein but not PIN2 protein ((Kharshiing et al. 2010) in the *pct1-2* mutant. If the short root *pct1-2* mutant is related to higher PAT (Al-Hammadi et al. 2003), why other auxin transport inhibitors like NPA are ineffective in rescuing this phenotype. It can be surmised that while our results indicate a putative linkage between GSH level and PAT in the *pct1-2* mutant, the linkage between these two responses can only be answered by determining the molecular identity of the gene causing *pct1-2* mutation. Our current efforts are directed towards the identification of the causative gene for *pct1-2* mutation by using emerging tools of genomics.

## Materials and methods

### Growth Conditions

The tomato (*Solanum lycopersicum*) genotype used in this study was a polycotyledon (*pct1-2*) mutant and its wild type in the Ailsa Craig (AC) background. Seeds were surface sterilized and germinated in the dark for approximately 3 days at 25±2ºC. After emergence of the radicle, seeds were transferred to 1/10 MS medium including 0.8% (w/v) agar with or without the supplement inhibitors of different concentration in a transparent plastic petri dish (9.5 cm *d* x 1.2 cm *h*). The germinating seedlings were subjected to chemical/inhibitors treatments after radical emergence from seed. Seedlings were grown under continuous white light (100 µmoles m^−2^s^−1^), and the hypocotyl and root length of seedlings were measured after 5-days of treatment. All experiments were independently repeated 3-4 times and for each set 10-15 seedlings were used (unless otherwise mentioned).

### Chemical treatments

In the chemical treatments, Glutathione (GSH); L-Buthionine Sulphoxime (BSO); brefeldin (BFA) (CAS: 20350-15-6); 2,3,5-Triiodo Benzoic acid (TIBA); 3-(5-[3, 4-dichlorophenyl]-2-furyl) acrylic acid (Gravacin); 2-(4-Diethylamino-2-hydroxybenzoyl) benzoic acid (BUM) (CAS: 5809-23-4), and 3-chloro-4-hydroxyphenylacetic acid (CHPAA) were used. All the chemicals were procured from Sigma-Aldrich, USA, except Gravacin (CAS: 188438-05-3) from ChemBridge, San Diego, CA, USA. The BUM was kindly provided by Prof. Markus Geisler (University of Zurich, Switzerland).

### Histochemical and Immunolocalization studies

For histochemical studies, 1 cm of primary root tips excised from 5-day old seedlings were fixed and hybridized as described by Riegler et al. (2008). Longitudinal sections (7µ) of root tips were used for the PIN1 immunolocalization studies. The immunolocalization was carried out as described by Kharshiing et al. (2010). The root sections were imaged using a laser-scanning microscope (Leica TCS SP2). Fluorescent labelled anti-PIN1 antibodies were excited with a laser beam at 488 nm, and emission was recorded. For all experiments, a minimum of five seedlings were analyzed, and at least three sections from each seedling were examined.

### *In-vivo* GSH Imaging by Monochlorobimane (MCB)

Monochlorobimane dyes are commonly used for GSH imaging. The stock solution of 100 mM MCB (Sigma Aldrich, USA) was prepared with dimethyl sulphoxide (DMSO). The stock solution was stored at −20°C. Aliquots were thawed immediately before use and diluted to final concentrations at 100 µM MCB. The samples were emerged to 150 µL of 100 µM MCB for 10 min and then washed with distilled water. The samples treated with 100 µM MCB were observed under a Leica Confocal microscope with excitation and emission wavelength of 385 nm and 485 nm respectively.

### Measurement of Total GSH

Total GSH was quantified in seed/seedling extracts by the spectrophotometry using glutathione reductase (GR) 5, 5’-dithio-bis (2-nitrobenzoic acid) (DTNB) recycling method, as described in Noctor and Foyer (1998). The seedlings were grown on 1/10 Murashige and Skoog media and were collected at different time points and snap frozen in liquid nitrogen. The frozen tissue was homogenized, extracted with 0.1 M HCl, and the supernatant was collected for GSH measurements. The assay was carried out in 120 mM NaH_2_PO4, 6 mM ethylenediaminetetraacetic acid (EDTA), 6 mM DTNB, 50 mM NADPH at 25°C, with 5-10 µL extract supernatant. The reaction was started by the addition of 0.5U GR (Sigma Aldrich) and monitored for 10 min by measuring A_412_ nm in a spectrophotometer. The GSH levels were calculated from A_412_ nm changes during the initial 5 min of reaction. Total glutathione levels were calculated using the equation of linear regression obtained from a standard GSH curve prepared in the same conditions as sample extracts.

### Microarray Analysis

Microarray analysis was carried out as described in details by Wang et al. (2009). Light-grown five-day-old *pct1-2* and WT seedlings were dissected to harvest to collect root, cotyledons, and hypocotyl. From harvested organs, the total RNA was isolated using the Qiagen RNeasy kit (Qiagen). The RNA was treated with DNase-treated as per manufacturer’s protocol (RNase-Free DNase Set; Qiagen). The transcripts level was checked by making a comparison of *pct1-2* and WT seedlings using two-color hybridizations onto EU-TOM1 microarray slides. For *pct1-2* mutant, those genes that displayed a log2 (ratio) higher than 0.5 or lower than −0.5 were considered as over- or under-expressed genes, which corresponds to >1.4.fold difference than WT.

## Acknowledgments

We thank Prof. Markus Geisler, Department of Plant and Microbial Biology, University of Zurich, CH-8008 Zurich, Switzerland for providing us BUM.

## Supplementary Data

**Supplementary Table S1.** Differentially Expressed Genes in different organ of *pct1-2* and WT seedlings.

**Supplementary Table S2**. Differentially Expressed Genes regulating GSH metabolism in different organ of *pct1-2* and WT seedlings.

